# Fast and robust imputation for miRNA expression data using constrained least squares

**DOI:** 10.1101/2021.11.03.467153

**Authors:** James W. Webber, Kevin M. Elias

## Abstract

High dimensional transcriptome profiling, whether through next generation sequencing techniques or high-throughput arrays, may result in scattered variables with missing data. Data imputation is a common strategy to maximize the inclusion of samples by using statistical techniques to fill in missing values. However, many data imputation methods are cumbersome and risk introduction of systematic bias. Here we present a new data imputation method using constrained least squares and algorithms from the inverse problems literature and present applications for this technique in miRNA expression analysis. The proposed technique is shown to offer an imputation orders of magnitude faster, with greater than or equal accuracy when compared to similar methods from the literature.

## 1 Background

Next generation sequencing technologies have revolutionized high-throughput analysis of the transcriptome. However, zero values present an inherent problem when analyzing the expression matrices generated through these techniques. When transcripts are relatively high in some samples, but not in others of the same type, or when the dimensionality of the data is high, technical zeros are even more likely to happen. Distinguishing technical zeros from true biologic null expression is essential for correct data interpretation.

To highlight the importance of data imputation, in terms of retaining classification accuracy, we consider the example problem of classifying images of handwritten digits. Classifying images of handwritten digits is a well studied problem in machine learning, and the test accuracy exceeds 99% using the state-of-the-art models [1]. See figure 1a, where we have shown an example, synthetic image of a handwritten 1. The same image, but with some missing pixels, is shown in figure 1b. The locations of the missing pixels are selected at random, and uniformly. If we wish to classify the images with missing pixels, then it is ill-advised to perform no data imputation (i.e., imputing zeros), as the accuracy would suffer. See figure 1c, where we have shown the effect of imputing zeros on the AUC, classification accuracy (ACC) and *F*_1_ score with the percentage of missing pixels. We see a decrease in classification accuracy and *F*_1_ score when more than 10% of the pixels are missing, and the reduction in accuracy is more pronounced as the percentage of missing pixels increases. Several strategies have been described for data imputation in gene expression and miRNA expression analysis [2, 3, 4, 5, 6, 7, 8, 9, 10, 11]. Two popular techniques are “VIPER” and “scImpute.” In [3], the authors introduce “VIPER”, which implements data imputation on gene expression data using a combination of lasso (or an elastic net), with a box constrained regression. That is, first a set of neighboring cells are found, which have related expression values to the missing cell, using lasso (or elastic nets). Then a box constrained regression is performed on the selected neighbors to fill in the missing gene expression values. The reason for using lasso as preprocessing, is that the quadratic programming code employed in [3] for box constrained regression does not scale well to large matrices, and thus lasso (or an elastic net) is used to select a subset of candidate nearest neighbors to reduce the array size before nonnegative regression. In [9], the authors introduce “scImpute”, which shares similarities in the intuition to VIPER. First, in the scImpute algorithm, the cells are clustered into *K* groups using spectral clustering [12]. Then, the missing cell expressions are reconstructed from their neighboring cells by a nonnegative constrained regression. That is, the missing values are imputed using nonnegative linear combinations (i.e., a linear combination with nonnegative coefficients) of their nearest neighbors, where the neighboring cells are determined by spectral clustering. We choose to focus on VIPER (specifically the lasso variant) and scImpute for comparison here, as they share the most similarities with the proposed method.

**Figure 1:**
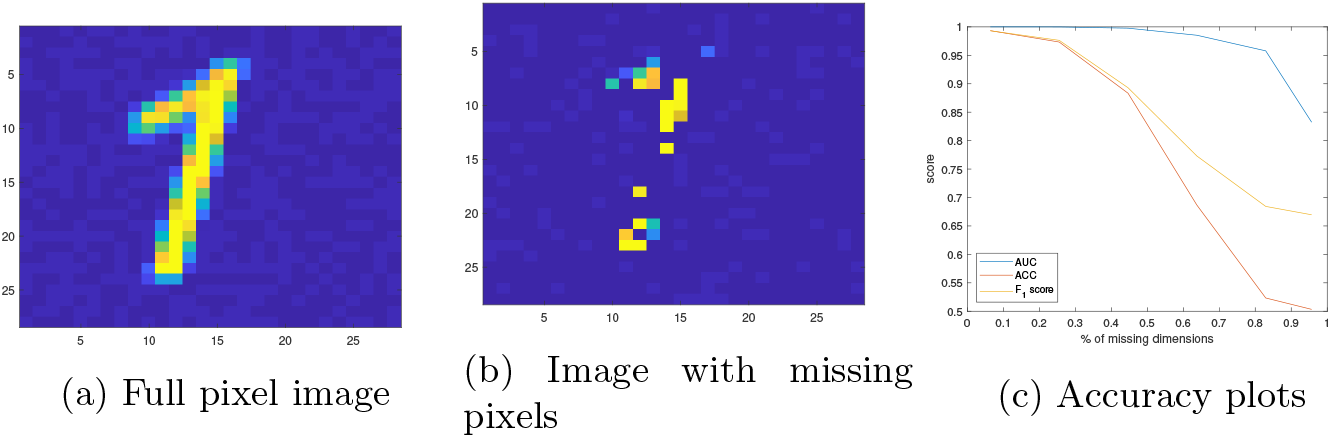
(A) - example image of a handwritten number 1. (B) - the same image but with missing pixels (C) - AUC, classification accuracy and *F*_1_ score with % of missing pixels.

Here we present a novel, fast method for data imputation using constrained Conjugate Gradient Least Squares (CGLS) borrowing ideas from the imaging and inverse problems literature. As an example of a desired application for this work, we present miRNA expression analysis, with a particular focus on cancer prediction. As shown below, highly accurate cancer prediction is possible using simple classifiers (e.g., a softmax function is used here), on a wide variety of data sets, in the case when all (or a large fraction) of the miRNA expression values are known, and there is little to no missing data. It is not always possible to measure all expression values contained in the training set, for every patient, however. To combat this, we aim to impute the missing values using the known expressions, so that we can retain use of our accurate model fit to the full set of miRNA available in the training set. We propose to reconstruct the missing data via nonnegative constrained regression, but with the further constraint that the regression weights sum to 1. Such constraints ensure that the imputed values lie within the range of the training data, with the idea to prevent overfitting. We enforce the regression weights to sum to 1 as a hard constraint in our objective, so that the nonnegative least squares and weight normalization steps are carried out simultaneously. To solve our objective, we apply the nonnegative Conjugate Gradient Least Squares (CGLS) algorithm of [13], typically applied in inverse problems and image reconstruction. The CGLS code we apply does not suffer the scaling issues encountered in, e.g., VIPER, for large matrices, and can process efficiently large scale expression arrays. The algorithm we propose offers a fast, efficient, and accurate imputation without the need for preprocessing steps, e.g., as in VIPER and scImpute. Our method is also completely nonparametric, and thus requires no tuning of hyperparameters before imputation, in contrast to scImpute and VIPER which require that two hyperparameters be tuned. Such parameters may be selected, for example, by cross validation, as is suggested in [3]. However, cross validation is slow, particularly for large data, and is thus impractical for clinical applications. To demonstrate the technique, we test the performance on miRNA expression data publicly available in the literature, and give a comparison to VIPER and scImpute. Specifically, as a measure of performance, we focus on how effectively each method retains the classification accuracy with the percentage of missing data (as in the curves shown in figure 1c). The proposed method is shown to be orders of magnitude faster than VIPER and scImpute, with greater than or equal accuracy, for the examples of interest considered here in cancer prediction.

## 2 Results

The method proposed here will be denoted by Fast Linear Imputation (FLI), for the remainder of this paper. The FLI algorithm and the core objective functions are discussed in detail in the appendix, section 5.1. In this section, we present a comparison of FLI, and the methods VIPER [3] and scImpute [9] from the literature on publicly available miRNA expression data [14, 15, 16] and synthetic handwritten image data. The specific implementations of VIPER and scImpute used here are discussed in the appendix, section 5.6. FLI is also compared against (unconstrained) regression, mean, and zero imputation as baseline. The classification model, selection of hyperparameters, and classification metrics are detailed in the appendix.

### 2.1 Synthetic handwritten image results

In this section, we present our results on the synthetic handwritten image data discussed in the introduction. For more details on this data see section 5.3. In figures 2a-2c we present plots of the AUC, ACC and *F*_1_ scores with the percentage of missing dimensions, for each method. Figures 2d-2f show the corresponding plots of the mean, standard deviation and maximum imputation errors, over all test patients. The plots in figures 2e and 2f are cropped on the vertical axis to better highlight the errors corresponding to the more competitive methods. This cuts off the end of error curve corresponding to scImpute, which spikes when ≈ 95% of the dimensions are missing. In table 1, we present the average values over the curves in figures 2a-2f, as a measure of the average performance over all possible levels of missing data. The mean, standard deviation, and maximum imputation times, over all test patients, are given in table 1c.

**Figure 2:**
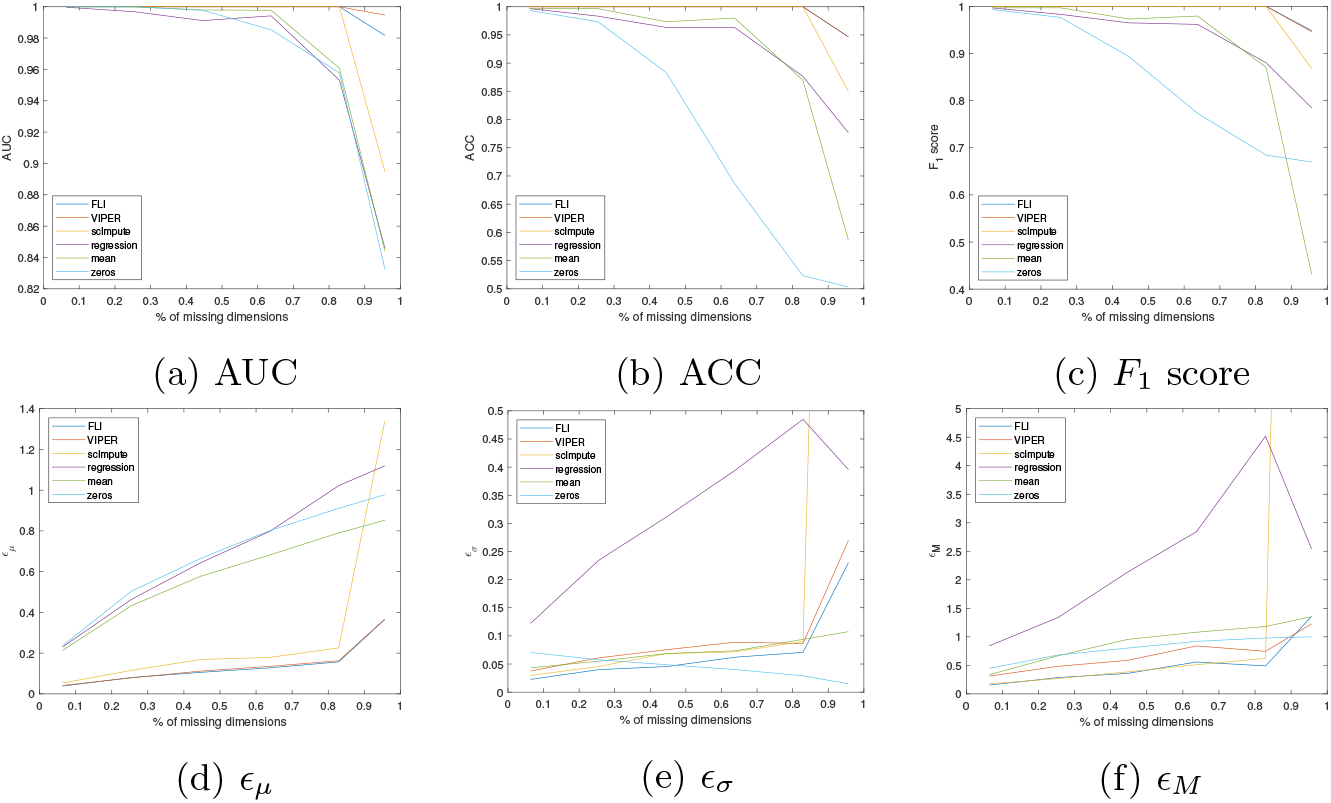
Handwritten image data results. (A)-(C) - AUC, ACC and *F*_1_ scores with percentage of missing dimensions. (D)-(F) - mean (*ϵ_μ_*), standard deviation (*ϵ_σ_*) and maximum (*ϵ_M_*) imputation errors over all test patients, with percentage of missing dimensions. The method is given in the figure legend.

**Table 1:** Handwritten image data results. (A) - mean values over curves shown in figures 2a-2c. (B) - mean values over curves shown in figures 2d-2f. (C) - mean (*t_μ_*), standard deviation (*t_σ_*), and maximum (*t*_max_) imputation times (in seconds) over all test patients. In table (A), ~ 1 indicates that the AUC is strictly greater than than .995. In table (C), ~ 0 indicates the imputation time is strictly less than .0005 seconds.

| Metric | FLI | VIPER | scImpute | regression | mean | zeros |
| --- | --- | --- | --- | --- | --- | --- |
| AUC | $\sim 1$ | $\sim 1$ | .98 | .96 | .97 | .96 |
| $F_1$ | .99 | .99 | .98 | .93 | .87 | .83 |
| ACC | .99 | .99 | .98 | .93 | .90 | .76 |

(a) Classification results
| Metric | FLI | VIPER | scImpute | regression | mean | zeros |
| --- | --- | --- | --- | --- | --- | --- |
| $\epsilon_\mu$ | .15 | .15 | .35 | .71 | .59 | .68 |
| $\epsilon_\sigma$ | .08 | .10 | .55 | .32 | .07 | .04 |
| $\epsilon_M$ | .53 | .70 | 6.2 | 2.4 | .93 | .81 |

(b) Imputation errors
| Time | FLI | VIPER | scImpute | regression | mean | zeros |
| --- | --- | --- | --- | --- | --- | --- |
| $t_\mu$ | .106 | 11.0 | 5.21 | .126 | $\sim 0$ | $\sim 0$ |
| $t_\sigma$ | .066 | 5.47 | 2.42 | .122 | $\sim 0$ | $\sim 0$ |
| $t_{\max}$ | .346 | 33.8 | 14.6 | 1.23 | $\sim 0$ | $\sim 0$ |

(c) Imputation time

In terms of retaining the classification accuracy, FLI, VIPER, and scImpute offer comparable performance. FLI and VIPER are joint best and offer mean AUC, ACC, and *F*_1_ scores exceeding 99%. For the baseline methods, namely (unconstrained) regression, mean, and zero imputation, we see a reduction in the classification accuracy, and the reduction is more pronounced when > 70% of the dimensions are missing, as evidenced by the curves in figures 2a-2c. In terms of the imputation error, FLI offers the most consistent imputation accuracy, when compared to VIPER and scImpute, in the sense that FLI offers the smallest standard deviation and maximum errors. For an example image imputation, see figure 3. where we have shown image reconstructions of the handwritten one image discussed in the introduction (figure 1a). We see ghosting artifacts in the VIPER reconstruction, and a significant blurring effect in the scImpute reconstruction. The regression imputation appears overfit, and introduces severe artifacts. FLI offers the clearest and sharpest image, with relatively few artifacts. So, in some cases, there are artifacts introduced by the VIPER and scImpute reconstructions. While this is not enough to confuse the classifier (i.e., the classification accuracy is still retained), the imputation error is less consistent when compared to FLI. In particular, the average maximum imputation error offered by FLI, over all levels of missing dimensions, is 17% lower than the next best performing method, namely VIPER. See the third row of table 1b. FLI is also orders of magnitude faster than VIPER and scImpute, as indicated by the imputation times of table 1c.

**Figure 3:**
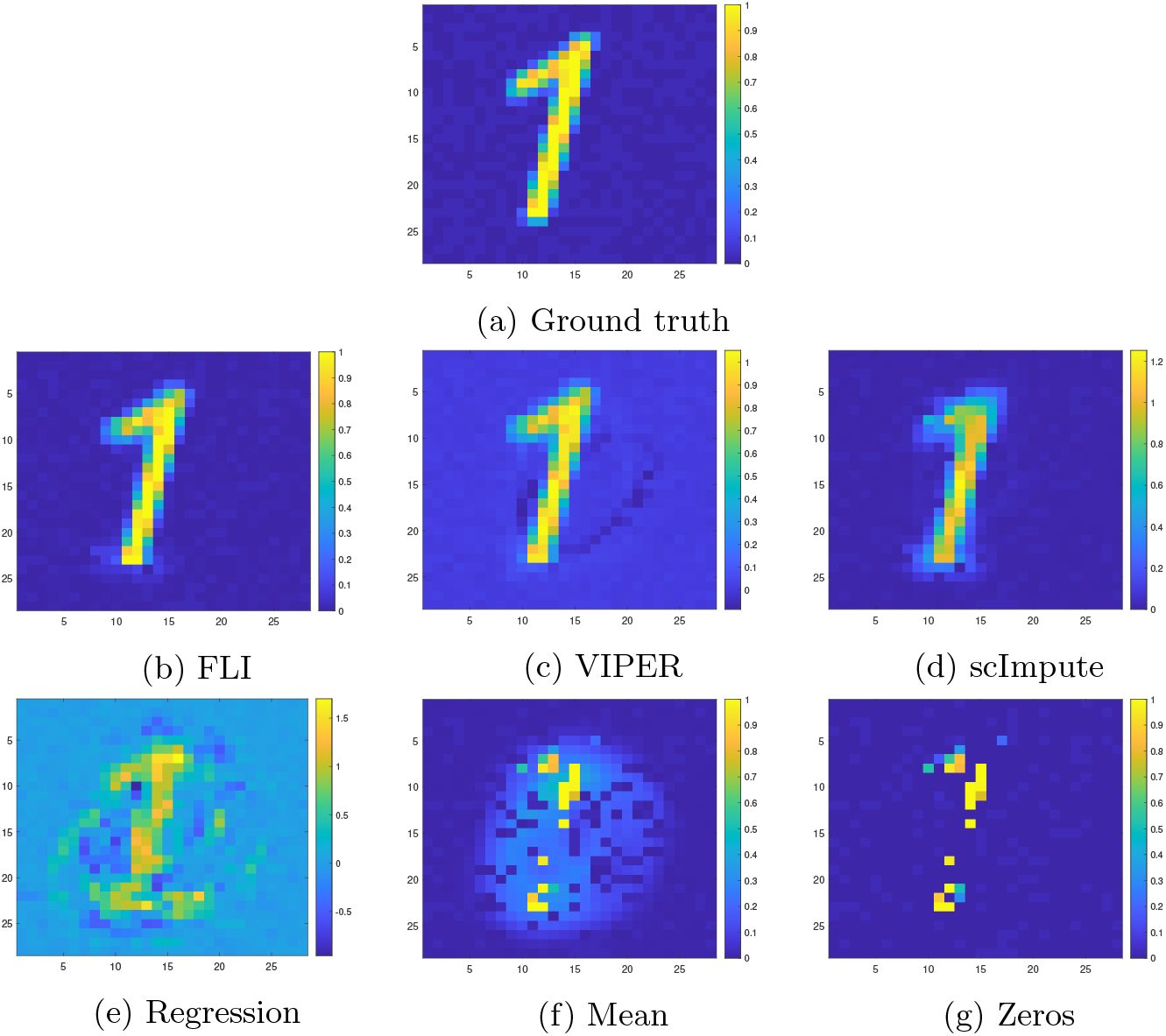
Example image reconstructions of the one image shown in the introduction, using all methods considered. The number of missing pixels is 550, which is 550/784 = 70% of all pixels. The ground truth is also shown for comparison.

### 2.2 Singapore results

Here we present our results on the miRNA expression data of Chan *et. al*. [16], collected from Singaporean patients. This data includes significant batch effects due to different measurement technologies. See section 5.3 for more details.

See figures 4a-4c for plots of the classification accuracy, and figures 4d-4f for the imputation errors with the percentage of missing dimensions. See table 2 for the mean values over the curves in figures 4a-4f, and the mean, standard deviation, and maximum imputation times. In this example, FLI offers the best performance in terms of retaining the AUC, ACC and *F*_1_ score, on average, across all levels of missing dimensions. As evidenced by figure 4a, FLI offers the highest AUC over all levels of missing dimensions. We see a similar effect in the ACC and *F*_1_ score curves of figures 4b and 4c, although, in a minority of cases, scImpute slightly outperforms FLI. The retention of the classification accuracy is significantly reduced using regression, mean and zero imputation, when compared to FLI, VIPER, and scImpute.

**Figure 4:**
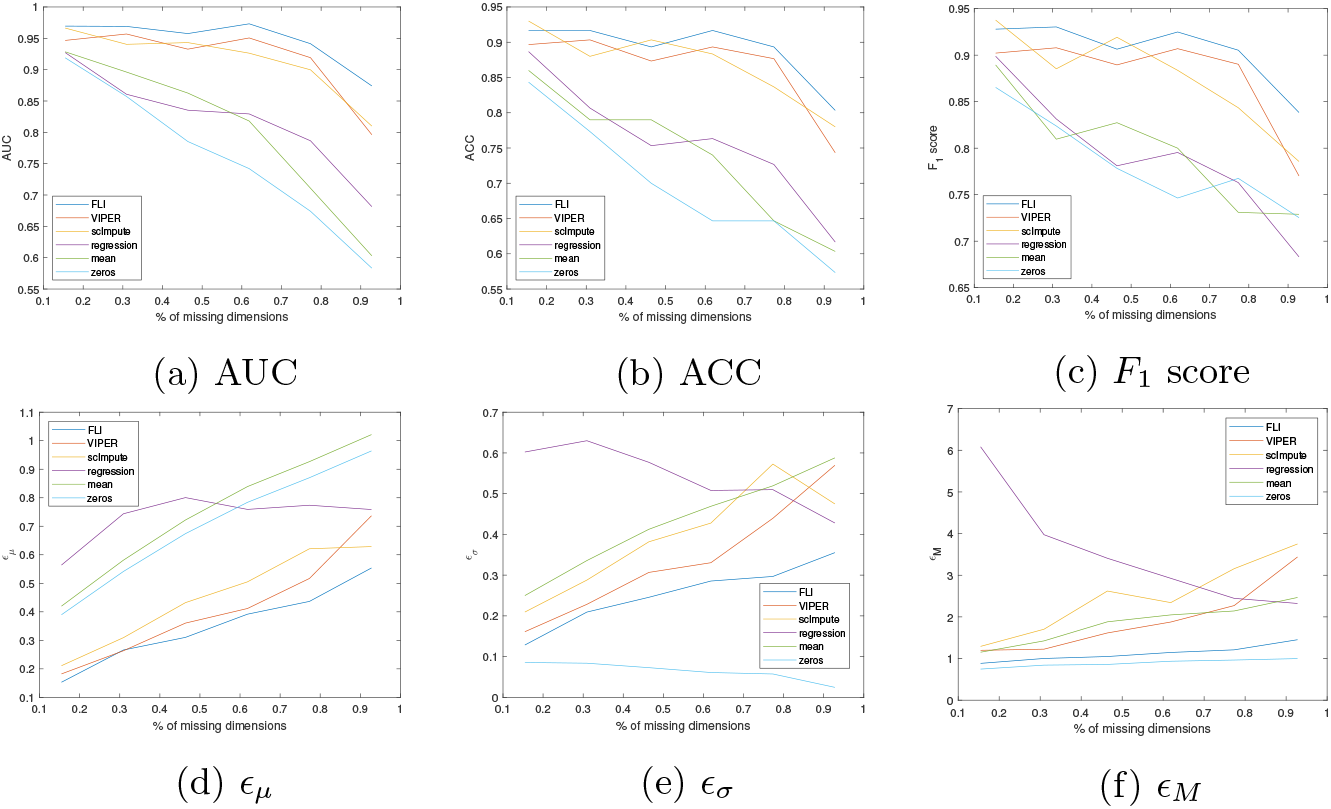
Singapore data results. (A)-(C) - AUC, ACC and *F*_1_ scores with percentage of missing dimensions. (D)-(F) - mean (*ϵ_μ_*), standard deviation (*ϵ_σ_*) and maximum (*ϵ_M_*) imputation errors over all test patients, with percentage of missing dimensions. The method is given in the figure legend.

**Table 2:** Singapore data results. (A) - mean values over curves shown in figures 4a-4c. (B) - mean values over curves shown in figures 4d-4f. (C) - mean (*t_μ_*), standard deviation (*t_σ_*), and maximum (*t*_max_) imputation times (in seconds) over all test patients. In table (C), ~ 0 indicates that the imputation time is strictly less than .0005 seconds.

| Metric | FLI | VIPER | scImpute | regression | mean | zeros |
| --- | --- | --- | --- | --- | --- | --- |
| AUC | .95 | .92 | .91 | .82 | .80 | .76 |
| $F_1$ | .91 | .88 | .88 | .79 | .80 | .78 |
| ACC | .89 | .86 | .87 | .76 | .74 | .70 |

(a) Classification results
| Metric | FLI | VIPER | scImpute | regression | mean | zeros |
| --- | --- | --- | --- | --- | --- | --- |
| $\epsilon_\mu$ | .35 | .41 | .45 | .73 | .75 | .70 |
| $\epsilon_\sigma$ | .25 | .34 | .39 | .54 | .43 | .06 |
| $\epsilon_M$ | 1.12 | 1.94 | 2.50 | 3.53 | 1.85 | .90 |

(b) Imputation errors
| Time | FLI | VIPER | scImpute | regression | mean | zeros |
| --- | --- | --- | --- | --- | --- | --- |
| $t_\mu$ | .009 | 7.29 | .100 | $\sim 0$ | $\sim 0$ | $\sim 0$ |
| $t_\sigma$ | .002 | 8.42 | .060 | $\sim 0$ | $\sim 0$ | $\sim 0$ |
| $t_{\max}$ | .027 | 49.2 | .300 | .003 | .001 | $\sim 0$ |

(c) Imputation time

FLI offers the most optimal performance in terms of the mean, standard deviation, and maximum imputation error, across all levels of missing dimensions, when compared to VIPER and scImpute. A zero imputation offers the best maximum and standard deviation error over all methods. The mean error offered by zero imputation is significantly higher than that of FLI, scImpute and VIPER, however. We would expect the standard deviation of a zero imputation to be low, as there is not much variation among the imputations (i.e., many of the values are zeros). The maximum error curves of figure 4f indicate that, for some patients, the imputation error is high when using FLI, VIPER and scImpute, as simply imputing zeros offers less error. Such erroneous patients can be considered outliers, and do not greatly effect the overall classification accuracy, as evidenced by the plots of figures 4a-4c.

The imputation time offered by FLI is orders of magnitude faster than VIPER and scImpute. For example, FLI is approximately three orders of magnitude faster, in terms of mean imputation time, when compared to VIPER, which was the next best performing method in terms of AUC, ACC and *F*_1_ score, after FLI. The imputation time offered by FLI is also more consistent when compared to VIPER, as evidenced by the *t_σ_* scores. When compared to scImpute, FLI is approximately one order magnitude faster in terms of mean and maximum imputation time, and is more consistent with lower standard deviation. Regression, mean and zero imputation are the fastest methods, but at the cost of accuracy.

This example was included given the presence of significant batch effects, as discussed at the beginning of this section, and in more detail in section 5.3. This example provides evidence that FLI is most optimal (compared to similar methods such as VIPER and scImpute), in terms of accuracy and imputation time, when imputing data in the presence of batch effects.

### 2.3 Korea results

Here we present our results on the miRNA expression data of Lee *et. al*. [17], collected from Korean patients. For more details on this data see section 5.3, point (3).

In figures 5a-5c we present plots of the AUC, ACC and *F*_1_ scores with the percentage of missing dimensions, for each method. Figures 2d-2f show the corresponding plots of the mean, standard deviation and maximum imputation errors. The plots in figures 5d and 5f are cropped to *E* = 0.35 on the vertical axis to better highlight the errors corresponding to the more competitive methods. Thus, parts of some of the error curves are missing in figures 5d and 5f. For example, in most cases (i.e., for most levels of missing dimensions considered), the zero imputation mean and maximum error exceeds 0.35 and thus why much of the light blue curves corresponding to zero imputation are missing in the plots. In tables 3a and 3b, we present the average values over the curves in figures 5a-5f. The mean, standard deviation, and maximum imputation times, over all test patients, are given in table 3c.

**Figure 5:**
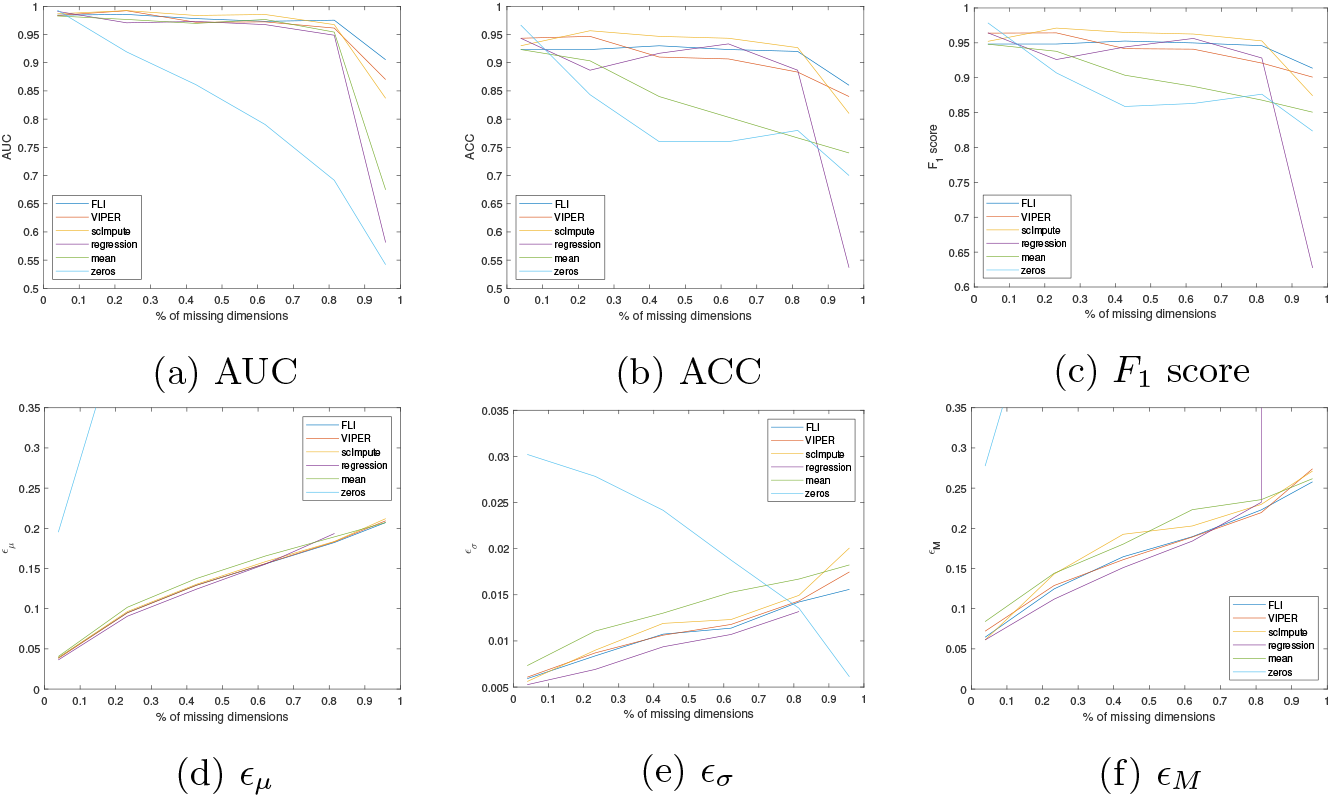
Korea data results. (A)-(C) - AUC, ACC and *F*_1_ scores with percentage of missing dimensions. (D)-(F) - mean (*ϵ_μ_*), standard deviation (*ϵ_σ_*) and maximum (*ϵ_M_*) imputation errors over all test patients, with percentage of missing dimensions. The method is given in the figure legend.

**Table 3:** Korea data results. (A) - mean values over curves shown in figures 5a-5c. (B) - mean values over curves shown in figures 5d-5f. (C) - mean (*t_μ_*), standard deviation (*t_σ_*), and maximum (*t*_max_) imputation times (in seconds) over all test patients. In table (C), ~ 0 indicates that the imputation time is strictly less than .0005 seconds.

| Metric | FLI | VIPER | scImpute | regression | mean | zeros |
| --- | --- | --- | --- | --- | --- | --- |
| AUC | .97 | .96 | .96 | .91 | .92 | .80 |
| $F_1$ | .94 | .94 | .95 | .89 | .90 | .88 |
| ACC | .91 | .91 | .92 | .85 | .83 | .80 |

(a) Classification results
| Metric | FLI | VIPER | scImpute | regression | mean | zeros |
| --- | --- | --- | --- | --- | --- | --- |
| $\epsilon_\mu$ | .13 | .13 | .14 | .27 | .14 | .67 |
| $\epsilon_\sigma$ | .01 | .01 | .01 | .17 | .01 | .02 |
| $\epsilon_M$ | .17 | .17 | .18 | 75.3 | .19 | .72 |

(b) Imputation errors
| Time | FLI | VIPER | scImpute | regression | mean | zeros |
| --- | --- | --- | --- | --- | --- | --- |
| $t_\mu$ | .024 | 18.2 | .365 | .005 | $\sim 0$ | $\sim 0$ |
| $t_\sigma$ | .016 | 5.33 | .348 | $\sim 0$ | $\sim 0$ | $\sim 0$ |
| $t_{\max}$ | .313 | 48.0 | 2.58 | .063 | $\sim 0$ | $\sim 0$ |

(c) Imputation time

**Table 4:** Keller and Japan data results. (A) - mean values over curves shown in figures 7a-7c. (B) - mean (*t_μ_*), standard deviation (*t_σ_*), and maximum (*t*_max_) imputation times (in seconds) over all test patients. In table (A), ~ 1 indicates that the AUC is strictly greater than .995.

| Classification | AUC | $F_1$ score | ACC |
| --- | --- | --- | --- |
| multiple sclerosis | .96 | .85 | .93 |
| melanoma | .97 | .88 | .93 |
| HCC | .99 | .93 | .97 |
| bladder cancer | $\sim 1$ | .97 | .99 |

(a) Classification results
| Classification | $t_\mu$ | $t_\sigma$ | $t_M$ |
| --- | --- | --- | --- |
| multiple sclerosis | .008 | .004 | .102 |
| melanoma | .009 | .006 | .128 |
| HCC | .273 | .155 | .922 |
| bladder cancer | .298 | .161 | 1.15 |

(b) Imputation times

In this example, FLI, VIPER, and scImpute offer similar levels of performance in terms of retaining the classification accuracy, with FLI offering the best average AUC, and scImpute the best average ACC and *F*_1_ scores. A standard regression imputations is also effective in retaining the classification accuracy up to approximately 85% of dimensions missing, and is comparable to FLI, VIPER, and scImpute within this range. We see a sharp reduction in accuracy in the regression curves (i.e., the purple curves of figures 5a-5c) when more than 85% of the dimensions are missing, however, and regression significantly underperforms FLI, scImpute, and VIPER at this limit. For mean and zero imputation, we see a more gradual reduction in accuracy, when compared to regression. The imputation errors offered by FLI, VIPER, scImpute, and mean imputation are comparable, and outperform regression and zero imputation. When compared to VIPER and scImpute, FLI offers an imputation time which is orders of magnitude faster, in terms of mean imputation time. The imputation time offered by FLI is also more consistent, with lower standard deviation and maximum, when compared to VIPER and scImpute. As was the case in the previous examples, regression, mean, and zero imputation are the fastest methods, but at the cost of accuracy.

## 3 Discussion

In this paper we introduced FLI, a fast, robust data imputation method based on constrained least squares. To illustrate the technique, we tested FLI on synthetic handwritten image and real miRNA expression data sets, and gave a comparison to two similar methods from the literature, namely VIPER [3] and scImpute [9]. We also compared against (unconstrained) regression, mean, and zero imputation as baseline. The results highlight the effectiveness of FLI in retaining the classification accuracy in cancer prediction applications using miRNA expression data, and in image classification. When compared to VIPER and scImpute, FLI was shown to offer greater than or equal imputation accuracy, with imputation speed orders of magnitude faster than scImpute and VIPER, in all examples considered. VIPER, scImpute, and FLI significantly outperformed regression, mean and zero imputation in terms of imputation accuracy, in all examples considered, but were slower given the greater computational complexity. For further validation of FLI on two more real miRNA expression data sets, see appendix 6.

In section 2.2, we considered an example expression data set collected from Singaporean patients, which included significant batch effects. When batch effects were present, FLI was shown to outperform VIPER and scImpute in terms of retaining the classification accuracy, and imputation error. On the handwritten image and Korean data sets, considered in sections 2.1 and 2.3, such batch effects were not detected. In these examples, the imputation accuracy offered by FLI, VIPER, and scImpute was comparable. This study provides evidence that FLI offers optimal imputation accuracy, when compared to the methods of literature, on batch data. This is important, since batch effects are common in medical data [18] and thus an imputation which is effective in combating batch effects, without the need for a-prioiri batch correction steps, is desirable.

In all examples considered, FLI was shown to be orders of magnitude faster than VIPER and scImpute. FLI is also completely nonparametric, and thus more straightforward to implement, in contrast to VIPER and scImpute, which require the tuning of two hyperparameters. It is suggested in [3] to tune the lasso parameter used by VIPER via cross validation. The reason for using lasso as preprocessing in VIPER, is that the quadratic programming code employed in [3] for nonnegative regression does not scale well to large matrices, and thus lasso (or elastic nets) are used to select a subset of candidate nearest neighbors before nonnegative regression.

A similar intuition is used in [9] in scImpute, whereby the training data is clustered into *K* groups using spectral clustering [12] before nonnegative regression. That is, the test samples are imputed using linear combinations of their nearest neighbors, where the neighbors are determined a-priori by spectral clustering. The algorithm we propose does not suffer such scaling issues for large matrices, and does not require any preprocessing steps before imputation. Our method is also completely nonparametric, and thus requires no tuning of hyperparameters before imputation, in contrast to scImpute and VIPER which require that two hyperparameters be tuned. Cross validation is slow, however, and resulted in long imputation times (in the order of minutes) when using VIPER. As noted by the authors in [3], the quadratic programming algorithm, used to implement box constrained regression, is slow, and thus why lasso preprocessing is proposed. The nonnegative least squares code of scImpute, applied in [9], also suffers efficiency issues. To combat this, the authors proposed to limit the number of regression weights a-priori using spectral clustering. FLI does not suffer such efficiency concerns, and requires no tuning of hyperparameters or preprocessing steps a-priori. FLI thus offers a faster and more straightforward imputation, when compared to VIPER and scImpute. This is important in applications where large numbers of samples need to be processed quickly (e.g., large gene expression arrays). In such applications, FLI offers the most practical imputation time, in comparison to VIPER and scImpute.

## 4 Conclusions and further work

The technique FLI proposed here offers accurate and fast imputation for miRNA expression data. In particular, the imputation offered by FLI was sufficiently accurate to retain the classification accuracy in cancer prediction and handwritten image recognition problems when a large proportion (up to 85%) of the dimensions were missing. Thus, FLI offers an effective means to classify samples with missing data, without the need for model retraining.

The application of focus here is miRNA expression analysis and cancer prediction. The current iteration of FLI requires a full training set (i.e., with no missing data) for the imputation, which can be considered a limitation of FLI. In further work we aim to address this limitation and develop FLI for more general missing data problems, and further apply FLI to other problems of interest in gene expression such as single-cell RNA seq. In this study, we assumed the locations of the missing data points to be random and uniform. In practice, the distribution of drop out events may be nonuniform. For example, in miRNA sequencing, the lower limit of detection is related to sequencing depth, thus within the technical zero range there may be a broad range of true expression values. We hypothesize that such expressions will more frequently be drop outs, when compared to more significantly expressed miRNA. In further work, we aim to test the effectiveness of FLI, and the methods of the literature, in the case when the drop out distribution is nonuniform, once the distribution of drop out events is decided upon.

## Appendix

### 5 Materials and methods

In this section we describe in detail our data imputation strategy FLI, and discuss the classification models and metrics used in our results.

#### 5.1 Description of FLI

Let 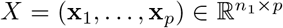 denote some training data, with no missing entries, and let 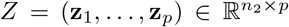 denote some unseen data with NaN columns. Let 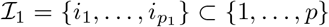 denote the indices of the NaN columns, and let 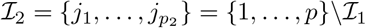, where *p* = *p*_1_ + *p*_2_. Then we aim to find

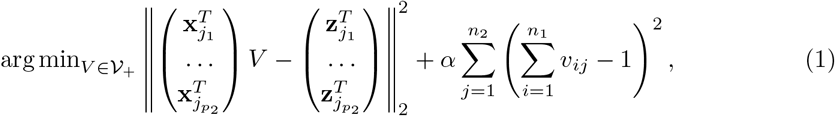

where

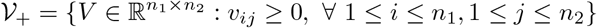

is the set of matrices size *n*_1_ × *n*_2_ with non-negative entries. Here *α* is set orders of magnitude larger than the maximum entry of *X* so that the columns of *V* all sum to 1 as a hard contraint. Specifically, we set *α* = 10^6^ × max_*i,j*_ (*X_ij_*). The missing values are then imputed via

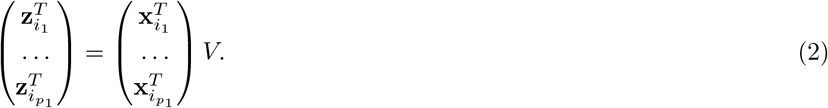

The *α* term and nonnegativity constraints ensure that the imputed values lie within the range of the training data. To solve (1) we apply the nonnegative CGLS algorithm of [13], typically applied in inverse problems and image reconstruction. Such imputation methods, which use linear combinations of the known expressions to reconstruct the missing values, are appropriate when there is a high level of linear dependence across expressions. The miRNA expressions are highly correlated, and thus FLI (and VIPER and scImpute) are appropriate for imputing miRNA expression data. See section 5.8, where we show singular value plots highlighting the linear dependence across micros for the data sets considered. The FLI algorithm detailed above is written in Matlab and is available from the authors upon request.

#### 5.2 Classification and error metrics

Here we introduce the metrics which will be used to assess the quality of the results. Let TP, FP, TN, and FN denote the number of true positives, false positives, true negatives, and false negatives, respectively, in a binary classification. Then, we report the following classification metrics in our comparisons:

1. The classification accuracy

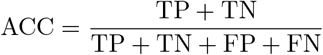
2. The *F*_1_ score

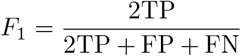
3. The Area Under the Curve (AUC) values corresponding to the Receiver Operator Characteristic (ROC).

All classification metrics above take values between 0 and 1. A value closer to 1 indicates a better performance, and vice versa.

Let **x**∈ ℝ^*p*^ denote a vector of ground truth expression values corresponding to a given patient, and let **x**_*ϵ*_ ∈ ℝ^*p*^ be an approximation (e.g., obtained through imputation). Then we define the least squares error

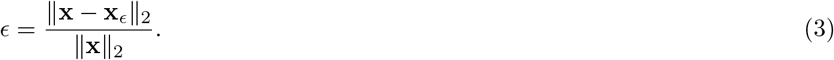

We use *ϵ* to measure the imputation error in section 2.

#### 5.3 Data sets

We consider the following four data sets from the literature for real data testing:

1. Serum miRNA expression data of [14, 15] collected from 2460 Japanese patients. Here the authors provide expression values for 1123 control patients, and for patients with 15 different types of diseases (the case number varies with the disease), including bladder cancer, hepatocellular carcinoma (HCC), breast cancer, ovarian cancer, and hepatitis. To compile the data used here, we combine the data sets of [14] and [15], making sure to delete any replicate patients. We focus on the HCC and bladder cancer case patients as most samples are provided for these diseases. We consider two separate binary classification problems, whereby we aim to separate the control set from bladder cancer and HCC patients.
2. The expression data of [19] collected by Keller *et. al*. from 454 German patients. The data is comprised of 70 healthy control patients, and patients with 14 different cancer and noncancer diseases (the case number varies with the disease), including melanoma, ovarian cancer, multiple sclerosis, and pancreatic cancer.
3. miRNA expression data of [17] collected by Lee *et. al*. from 232 Korean patients. This data is comprised of 88 patients with Pancreatic Cancer (PC), and 19 healthy controls. In [17], the authors combine the 19 healthy patients with 10 cholelithiasis patients to form a larger control set of 29 patients for use in the PC classifications. We use the same control set here, and aim to separate the controls from the PC patients. In total we consider *n* = 117 patients. The authors provide *p* = 2578 expression values, for each patient, all of which will be used in our classifications.
4. miRNA expression data of [16] collected by Chan *et. al*. from 116 Singaporean patients, consisting of 67 patients with breast cancer and 49 healthy controls. There are *p* = 116 expression values for each patient. This data is comprised of two cohorts, one size *n* = 62, and the other *n* = 54. The expression values of each cohort are measured using different technologies, which creates a batch effect. This example is included to test the effectiveness of each considered imputation method in the presence of batch effects.
5. Synthetic handwritten image data - the “Digits” database from Matlab. We focus on the images of zeros and ones provided for our classifications. That is, we consider a binary classification problem whereby we aim to separate images of zeros from images of ones. The data consists of 988 images of zeros and 1026 images of ones, of size 28 × 28. In this case, *n* = 1026 + 988 = 2014, and *p* = 28^2^ = 784.

#### 5.4 Classification model

Here we discuss the classification model used to classify patients in the real miRNA expression data experiments conducted in this paper. Let *X* ∈ ℝ^*n*×*p*^ be a set of imputed miRNA expression data (i.e., with no missing values). To classify patients, we train a softmax function [20] classification model

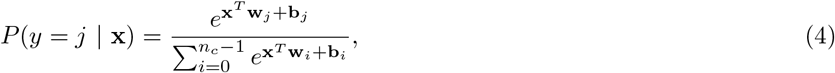

where *j* ∈ {0, 1*,…, n_c_* − 1} is the class label, with *n_c_* > 1 the number of classes, **x** ∈ ℝ^*p*^ is a patient sample after imputation (e.g., one row of *X*), and the (**w**_*j*_, **b**_*j*_) are weights and biases to be trained. Here *y* denotes the class label assigned to **x**. The class with the highest probability *P* is then chosen for membership.

#### 5.5 Validation methods

Here we discuss the validation methods used in the experiments conducted. We use a multiple hold out set validation. That is, we randomly and uniformly set aside a subset of samples size *n_t_ < n*_1_ from the data. After which, we randomly and uniformly assign a subset of *m < p* variables to NaN for each patient, where *m* is determined by the percentage of missing dimensions (see section 2). Then the missing dimensions are imputed, and the test patients are classified using the model discussed in section 4.The above process is repeated over *n_T_* trials and the results are averaged. For the data sets with small *n*, i.e., data sets (2)-(3) of section 5.3, where *n* < 150, we set *n_t_* = 10 and *n_T_* = 30. For data set (1) of section 5.3, where *n* > 1000, we set *n_t_* = 300 and *n_T_* = 1. In total there are *n_T_* × *n_t_* = 300 test trials for each classification performed.

#### 5.6 Implementation of VIPER and scImpute

Here we discuss the implementations of VIPER and scImpute used in this paper. Both algorithms are implemented in Matlab. The R code provided in [3, 9] was ran from within Matlab. The Matlab formulations of VIPER and scImpute are available from the authors upon request.

##### 5.6.1 scImpute

We perform scImpute as explained in [9], but with a few technical changes which we shall now discuss. First, since no outliers were detected in the data sets used here, we do not implement the outlier removal stage of scImpute discussed in [9]. In some instances, the Nonnegative Least Squares (NNLS) solver employed in [9] suffered crashing issues for underdetermined systems. Hence, in the case of undermined system matrices, we multiplied both sides by the matrix transpose so that the input to the NNLS solver was a square matrix. That is, we solved the normal equations, which is an equivalent problem. In rare cases, the NNLS solver produced the error “Matrix inversion failed”. In such instances, we imputed the mean value over the nearest neighbors determined by spectral clustering (i.e., the spectral clustering step described in [9]).

##### 5.6.2 VIPER

VIPER is implemented exactly as discussed in [3], with lasso used as preprocessing.

In [3, 9], the authors propose methods to determine the locations of the missing data. In this paper, we assume knowledge of the locations of the missing data a-priori, and hence we do not implement such aspects of VIPER and scImpute.

#### 5.7 Hyperparameter selection

Here we discuss the selection of hyperparameters for the methods compared against. For scImpute, there are two hyperparameters, namely the number of clusters *K* and the affinity parameter *σ* used for the spectral clustering step. Here we are using the notation of [9] and [12]. In [12], it is suggested to choose *σ* so that the within cluster variances are minimized. We use a similar idea and choose the *K* ∈ {2*,…*, 10} and *σ* such that the ratio of the between and within cluster variances is maximized, i.e., we maximize the Calinski-Harabasz statistic [21]. In [9], no method for choosing *K* or *σ* is given so we use the ideas of [12] (cited in [9]) to choose *K* and *σ*.

For lasso VIPER, the lasso parameter is chosen by 10-fold cross validation, and the threshold parameter is set to *t* = 0.001, as in [3]. Note here we are using the notation of [3].

As discussed in section 5.1, the *α* parameter of FLI (see equation (1)) is set orders of magnitude larger than the maximum entry of the expression training matrix *X*, so that the imputation weights all sum to 1 as a hard constraint. Specifically, *α* = 10^6^ × max_*i,j*_ (*X_ij_*) is set six orders of magnitude larger than the maximum entry of *X*.

The remaining methods compared against for baseline, namely regression, mean and zero imputation, have no hyperparameters.

#### 5.8 Singular value plots

Here we show plots of the singular values of the expression matrices *X* corresponding to some of data sets considered. See figure 6. We see a high level of linear dependence among the expression values, as indicated by the singular value plots. The nuclear norm [22] is commonly used to approximate the matrix rank. We use the nuclear norm here to measure the level of linear dependence among the miRNA expression values. A smaller nuclear norm indicates greater linear dependence, and vice-versa. Based on the nuclear norm values, there is higher linear dependence among the Japanese patients, and thus an imputation using linear combinations (e.g., as with FLI) is likely to be more accurate. The Keller data shows the least linear dependence among the expression values, and thus we expect the FLI imputation (and those of VIPER and scImpute) to be less accurate in this case.

**Figure 6:**
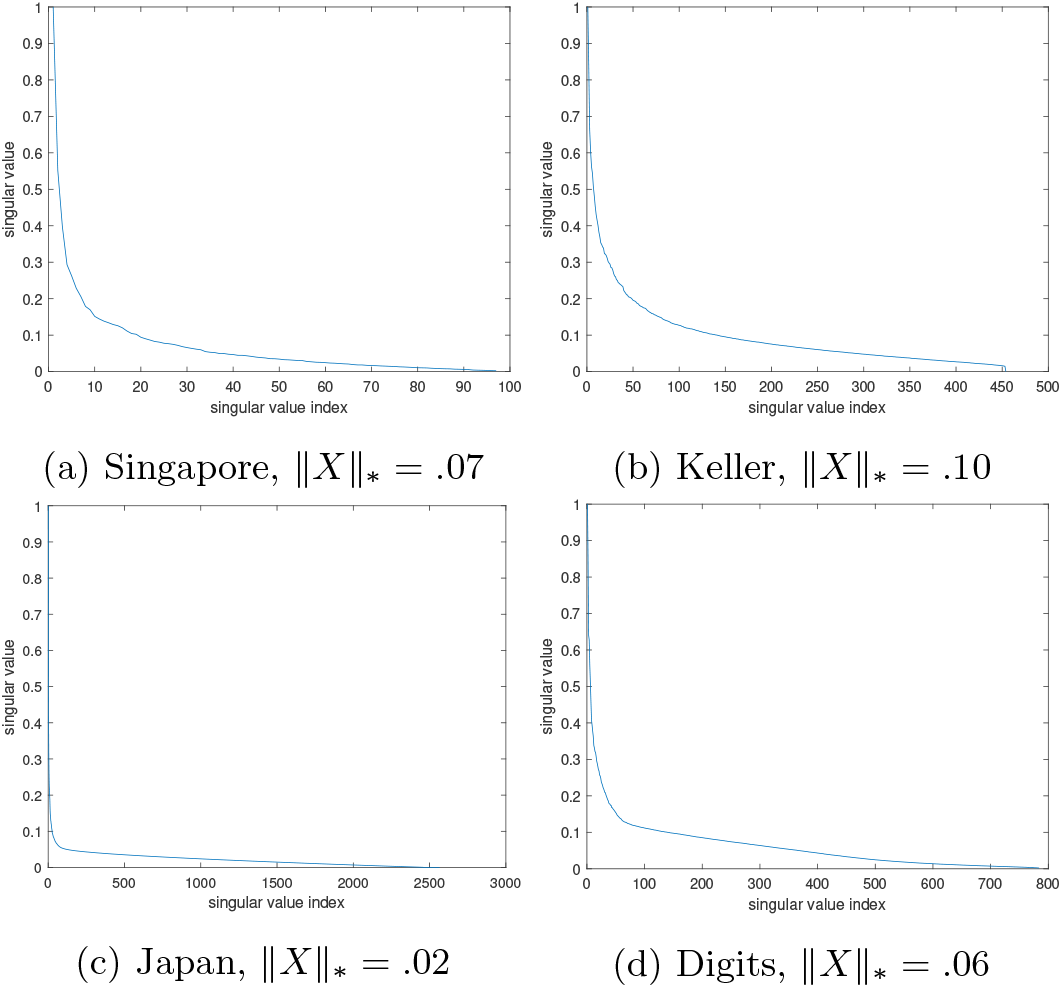
Singular value plots of the expression matrices *X* corresponding to the Japanese, Singapore, handwritten digits, and Keller data. The nuclear norm values ∥*X*∥_*_ are given in the figure subcaptions.

### 6 Additional results

Here we present additional validation of FLI on two more real miRNA expression data sets from the literature.

#### 6.1 Japan and Keller data results

In this section, we test the accuracy of FLI on the miRNA expression data of [14, 15], collected from Japanese patients, and the expression data of Keller *et. al*. [19], collected from German patients. For the Japanese data, we focus specifically on the binary classification problems of separating bladder cancer and HCC patients from controls, as is considered in [14, 15]. For the Keller, we focus on separating melanoma and multiple sclerosis patients from the control set. We chose melonoma and multiple sclerosis, as the softmax function classifier (discussed in section 4) offered the best accuracy scores in terms of AUC when separating these diseases from controls, compared to the other diseases considered by Keller. In total, we consider four binary classification problems, two associated with the Japanese data (i.e., the HCC and bladder cancer classifications), and two from the Keller data set (i.e., the melanoma and multiple sclerosis classifications). See figure 7 where we show curves of the classification accuracy scores with the percentage of missing dimensions, for each of the four classifications considered. In table 4 we present the average scores over the curves shown in figure 7, as an average measure of the performance of FLI over all levels of missing dimensions.

**Figure 7:**
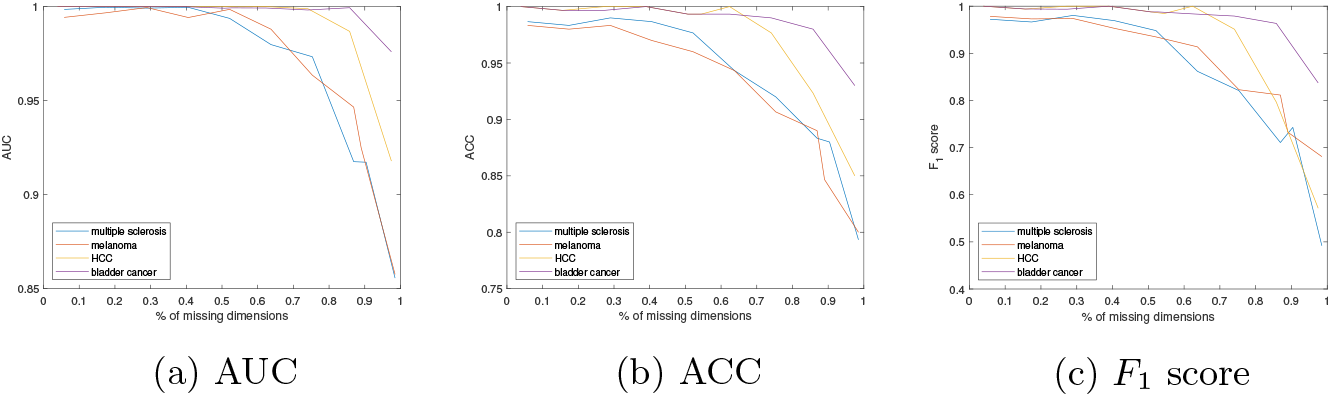
Keller and Japan data results, using FLI. (A)-(C) - AUC, ACC and *F*_1_ scores with percentage of missing dimensions. The disease classified is given in the figure legend.

The results offer further validation of the effectiveness of FLI in retaining the classification accuracy on two more large miRNA expression data sets from the literature. The imputation times of table 4b further validate the efficiency of FLI. For example, the maximum imputation time recorded, over all test patients processed, was *t_M_* = 1.15s. On the Japanese data, FLI is most effective in retaining the classification accuracy, when compared to the Keller data, and offers AUC, ACC, and *F*_1_ scores exceeding .95 up to 70% of dimensions missing. On the Keller data, FLI is less effective in retaining the classification accuracy, and offers AUC, ACC, and *F*_1_ scores exceeding .95 up to approximately 50% of dimensions missing. The mean curve values presented in table 4a highlight the difference in the effectiveness of FLI on the Japanese and Keller data sets. In figure 6, we showed singular value plots of the expression data matrices corresponding to the Keller and Japanese data sets. The plots indicated a higher level of linear dependence among the Japanese miRNA set, when compared to that of Kelller. Thus we would expect an imputation such as FLI, which uses linear combinations of the training expressions to impute the missing expressions, to be less effective on the Keller data set, as there is less linear dependence among the miRNA subset chosen by Keller. The results observed here are thus in line with the findings of figure 6.

## Abbreviations

FLI: Fast Linear Imputation
CGLS: Conjugate Gradient Least Squares

## Availability of data and materials

All real miRNA expression data sets considered here are publicly available online. See section 5.3for more details. The synthetic handwritten digit data is available for Matlab subscribers. The FLI code introduced here is available from the authors upon reasonable request.

## Ethics approval and consent to participate

No new human or animal data is presented here.

## Consent for publication

There are no issues regarding consent for publication.

## Authors’ contributions

JWW developed FLI and the original idea. The experiments were conducted by JWW. Analyses of results by JWW and KE. KE was as major contributor in writing the manuscript, and provided expert insight from a medical background needed to communicate this work to a medical audience.

